## Supplementary Figures for "A lineage tree-based hidden Markov model to quantify cellular heterogeneity and plasticity"

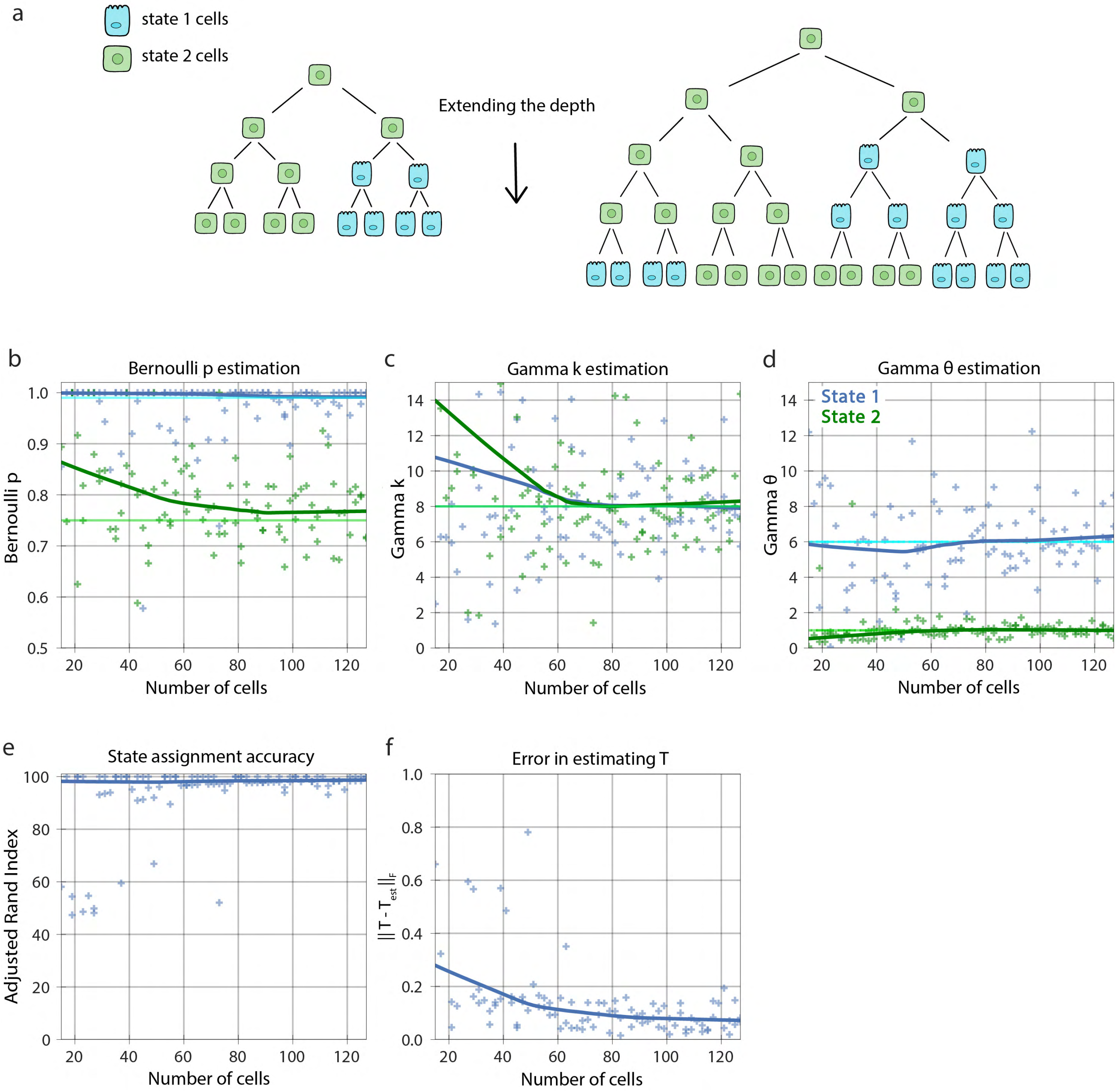


**Supplementary Figure 1: Performance on synthetic uncensored single lineages of increasing size with two states.** **(a)** Visual representation of increasing the lineage size with two states. **(b)** The Bernoulli parameter for states 1 and 2 as the number of cells increase. **(c)** The shape parameter, k, and **(d)** scale parameter, θ, of the Gamma distribution corresponding to the cell lifetime for states 1 and 2 as the number of cells increase. **(e)** The state assignment accuracy as the number of cells increases. **(f)** The error in the estimate of the transition probability matrix, T as the number of cells increase. In **(b-d)** the light green and blue solid lines show the true value of the parameters, and the dark green and blue solid lines show the Lowess trend of estimations.

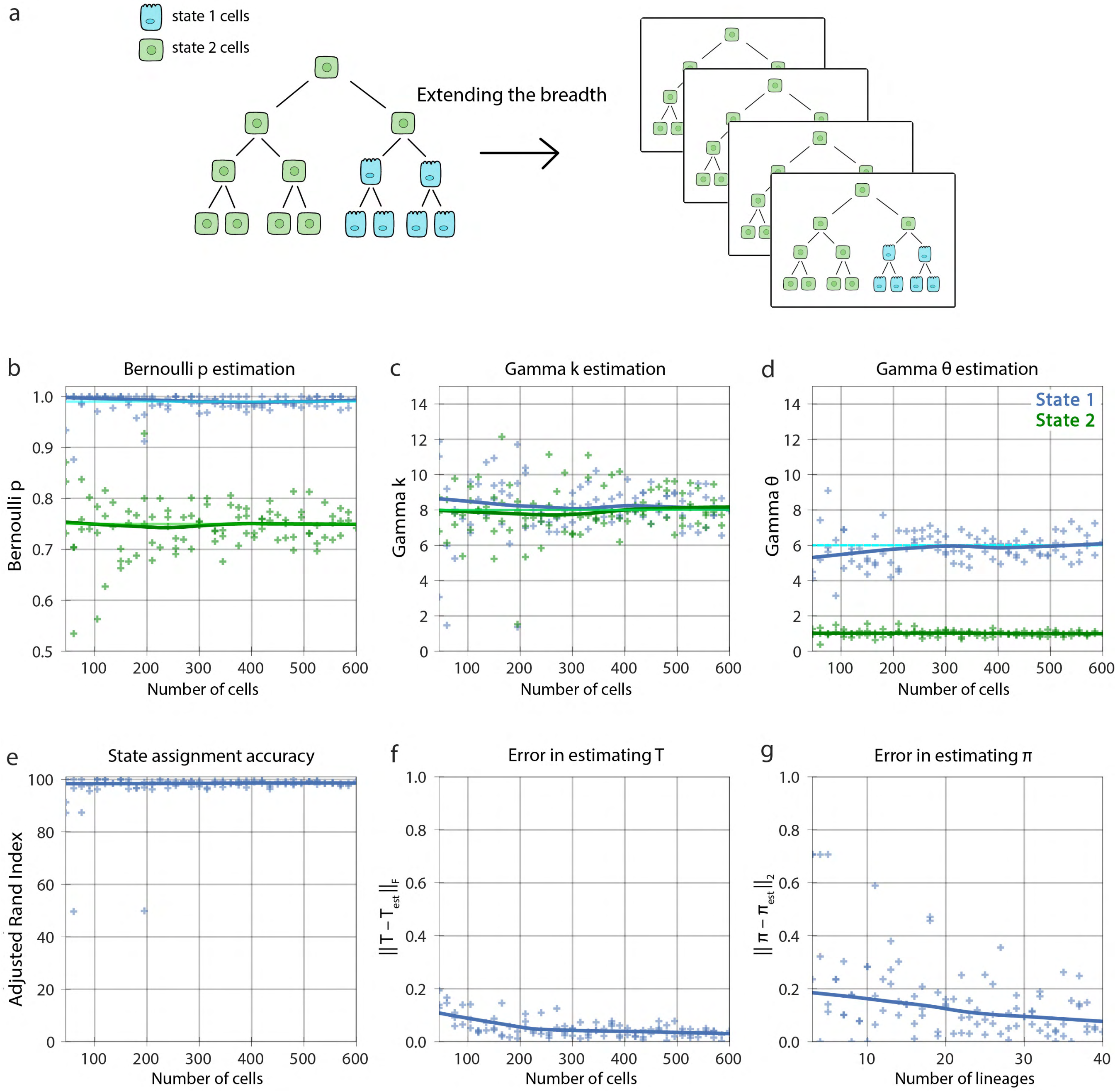


**Supplementary Figure 2: Performance on synthetic uncensored lineages of increasing number with two states. (a)** Visualization of increasing the number of fully observed lineages. **(b)** The Bernoulli parameter for states 1 and 2 as the number of cells increase. **(c)** The shape parameter, k, and **(d)** scale parameter, θ, of the Gamma distribution corresponding to the cell lifetime for states 1 and 2 as the number of cells increase. **(e)** The state assignment accuracy as the number of cells increases. **(f)** The error in the estimate of the transition probability matrix, T, as the number of cells increase. **(g)** The errors in the estimate of the initial probability matrix, π, as the number of lineages increase. In **(b-d)** the light green and blue solid lines show the true value of the parameters, and the dark green and blue solid lines show the Lowess trend of estimations.

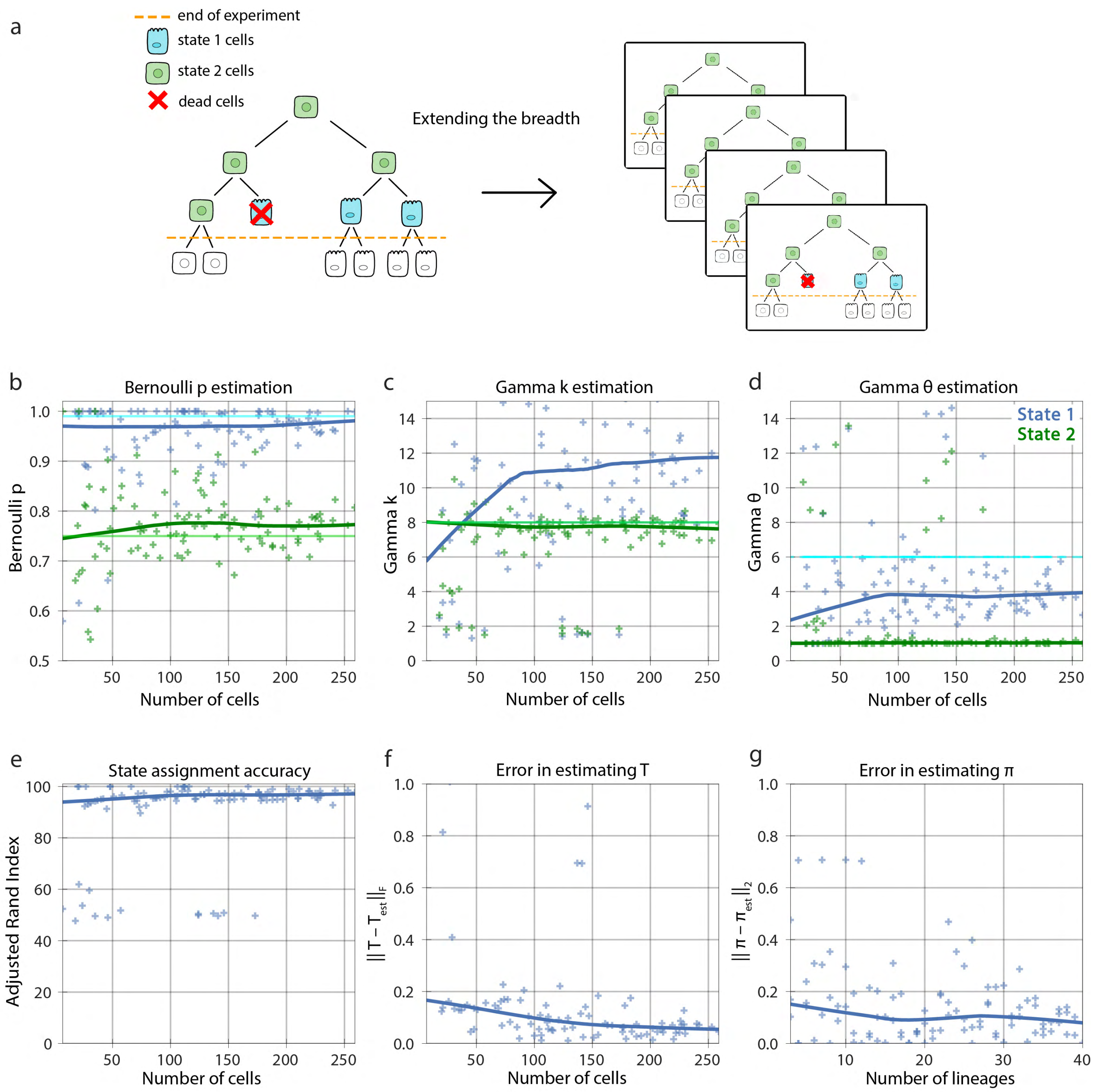


**Supplementary Figure 3: Performance on synthetic censored lineages of increasing number with two states. (a)** Visualization of the number of censored lineages increasing. **(b)** The Bernoulli parameter for states 1 and 2 as the number of cells increase. **(c)** The shape parameter, k, and **(d)** scale parameter, θ, of the Gamma distribution corresponding to the cell lifetime for states 1 and 2 as the number of cells increase. **(e)** State assignment accuracy as the number of cells increase. **(f)** The error in the estimate of the transition probability matrix, T, as the number of cells increase. **(g)** The errors in the estimate of the initial probability matrix, π, as the number of lineages increase. In **(b-d)** the light green and blue solid lines show the true value of the parameters, and the dark green and blue solid lines show the Lowess trend of estimations.

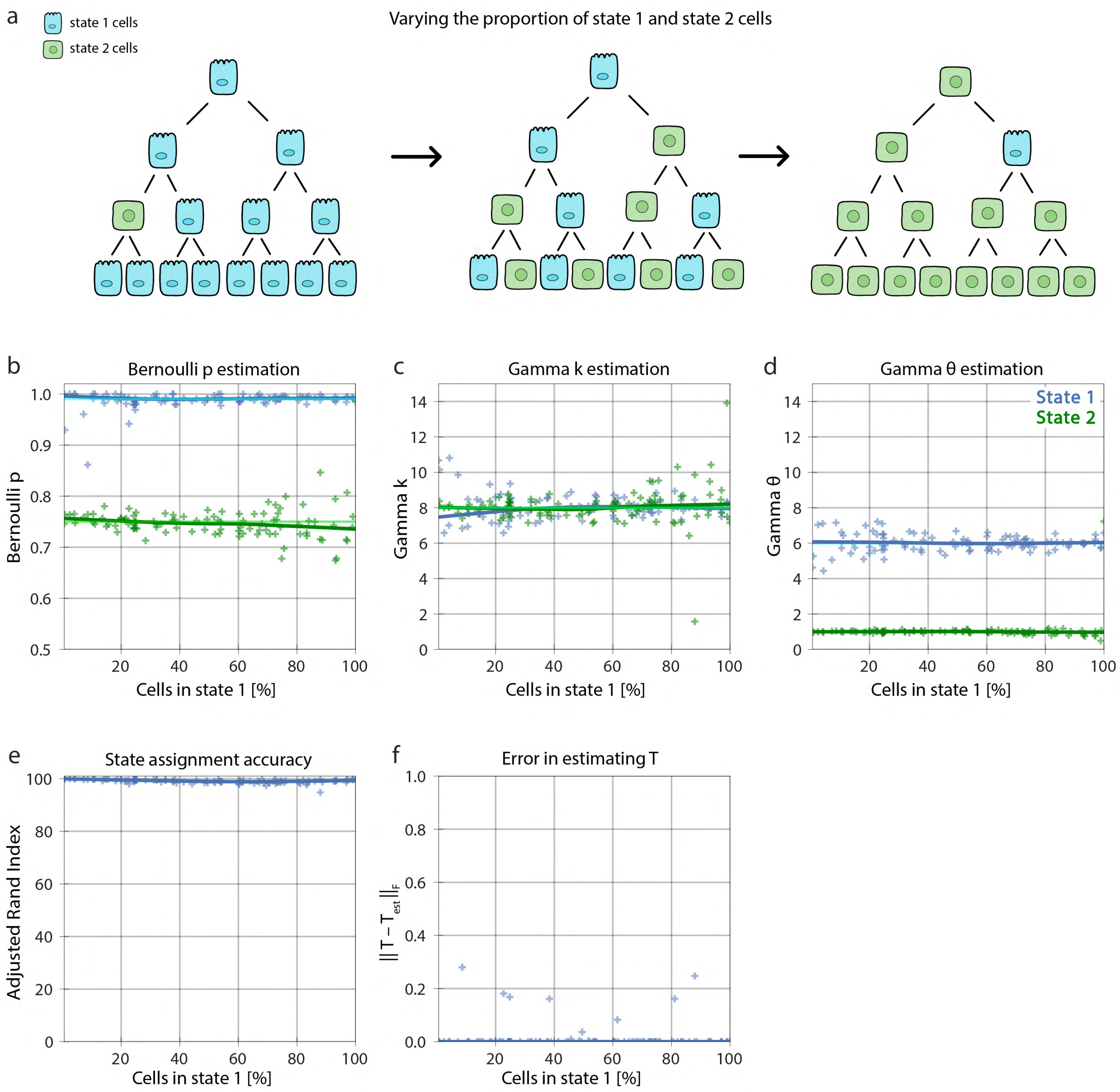


**Supplementary Figure 4: Model performance relative to the presence of each state for an uncensored lineage in a synthetic two-state dataset. (a)** Visualization of the distribution of cells in the lineage transitioning between state 1 and state 2. **(b)** The Bernoulli parameter for states 1 and 2 as the proportion of cells in state 1 increase. **(c)** The shape parameter, k, and **(d)** scale parameter, θ, of the Gamma distribution corresponding to the cell lifetime for states 1 and 2 as the proportion of cells in state 1 increase. **(e)** The state assignment accuracy as the proportion of cells in state 1 increase. **(f)** The errors in the estimate of the transition probability matrix, T, as proportion of cells in state 1 increase. In **(b-d)** the light green and blue solid lines show the true value of the parameters, and the dark green and blue solid lines show the Lowess trend of estimations.

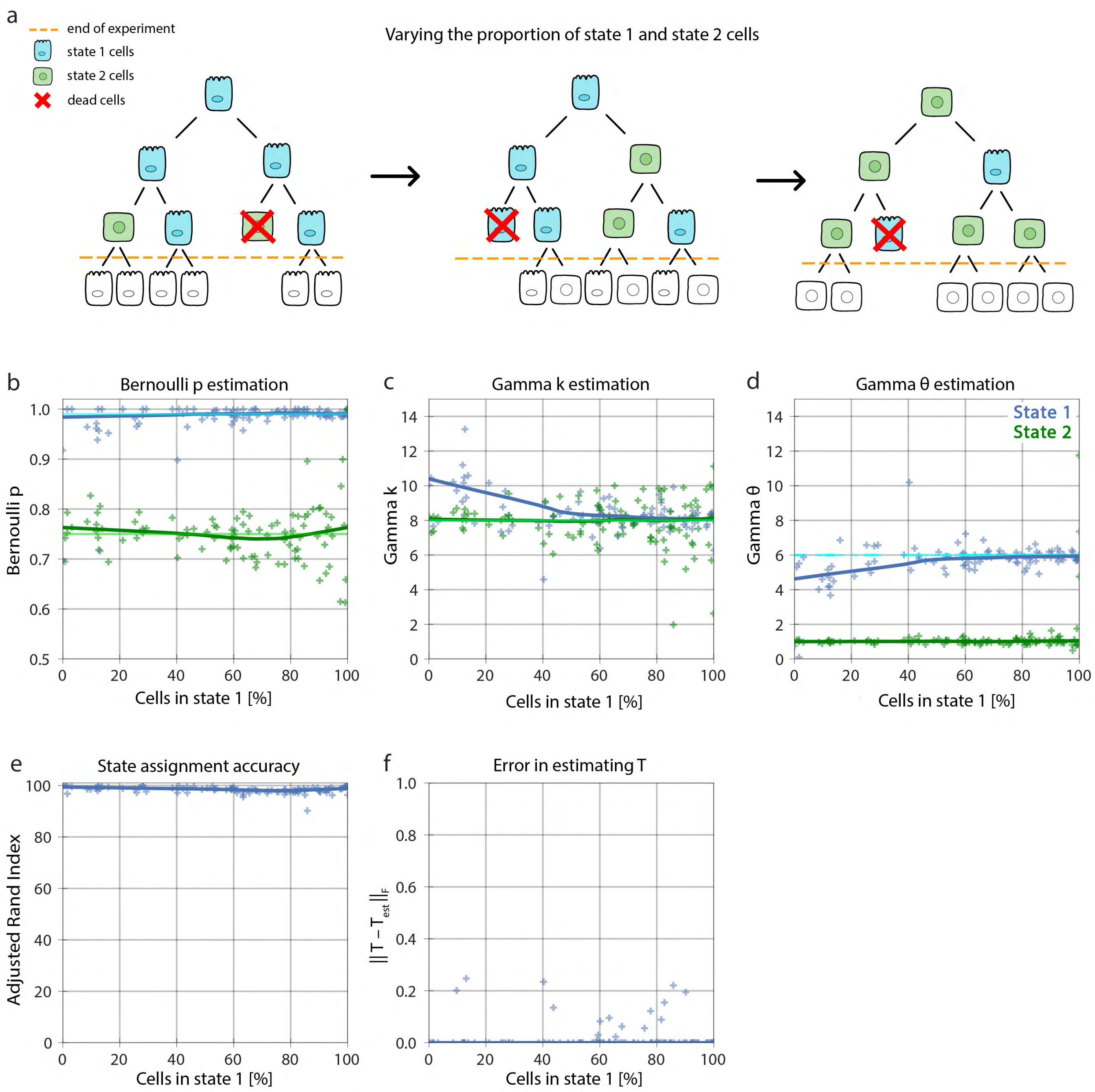


**Supplementary Figure 5: Change in model performance when varying the presence of a state for a censored lineage in a synthetic two-state dataset. (a)** Visualization of the proportion of cells in a censored lineage transitioning between state 1 and 2. **(b)** The Bernoulli parameter for states 1 and 2 as the proportion of cells in state 1 increase. **(c-d)** The shape parameter, k, and scale parameter, θ, of the Gamma distribution corresponding to the cell lifetime for states 1 and 2 as the proportion of cells in state 1 increase. **(e)** The state assignment accuracy as the proportion of cells in state 1 increase. **(f)** The error in the estimate of the transition probability matrix, T, as the proportion of cells in state 1 increase. In **(b-d)** the light green and blue solid lines show the true value of the parameters, and the dark green and blue solid lines show the Lowess trend of estimations.

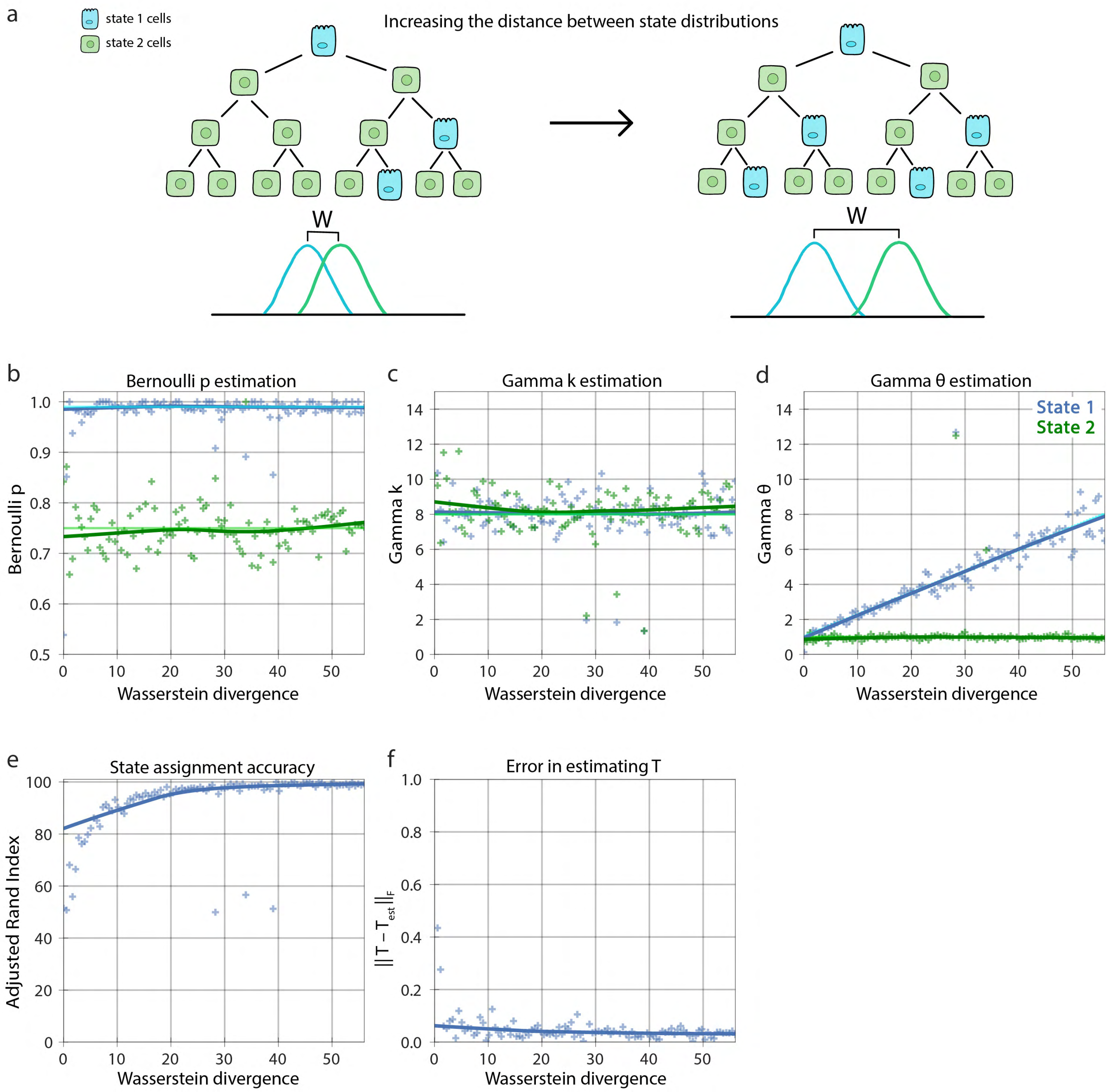


**Supplementary Figure 6: Change in model performance when varying state distribution similarity for an uncensored population of lineages in a synthetic two-state dataset. (a)** Visualization of the Wasserstein divergence increasing as the state distribution in the lineage varies. **(b)** The cell fate Bernoulli parameter compared to the true value. **(c)** The shape parameter, k, **(d)** the scale parameter, θ, of the Gamma distribution corresponding to the cell lifetime compared to the true values. **(e)** The state assignment accuracy as the Wasserstein divergence increases. **(f)** The errors in the estimate of the transition probability matrix, T, as the Wasserstein divergence increases. In **(b-d)** the light green and blue solid lines show the true value of the parameters, and the dark green and blue solid lines show the Lowess trend of estimations.

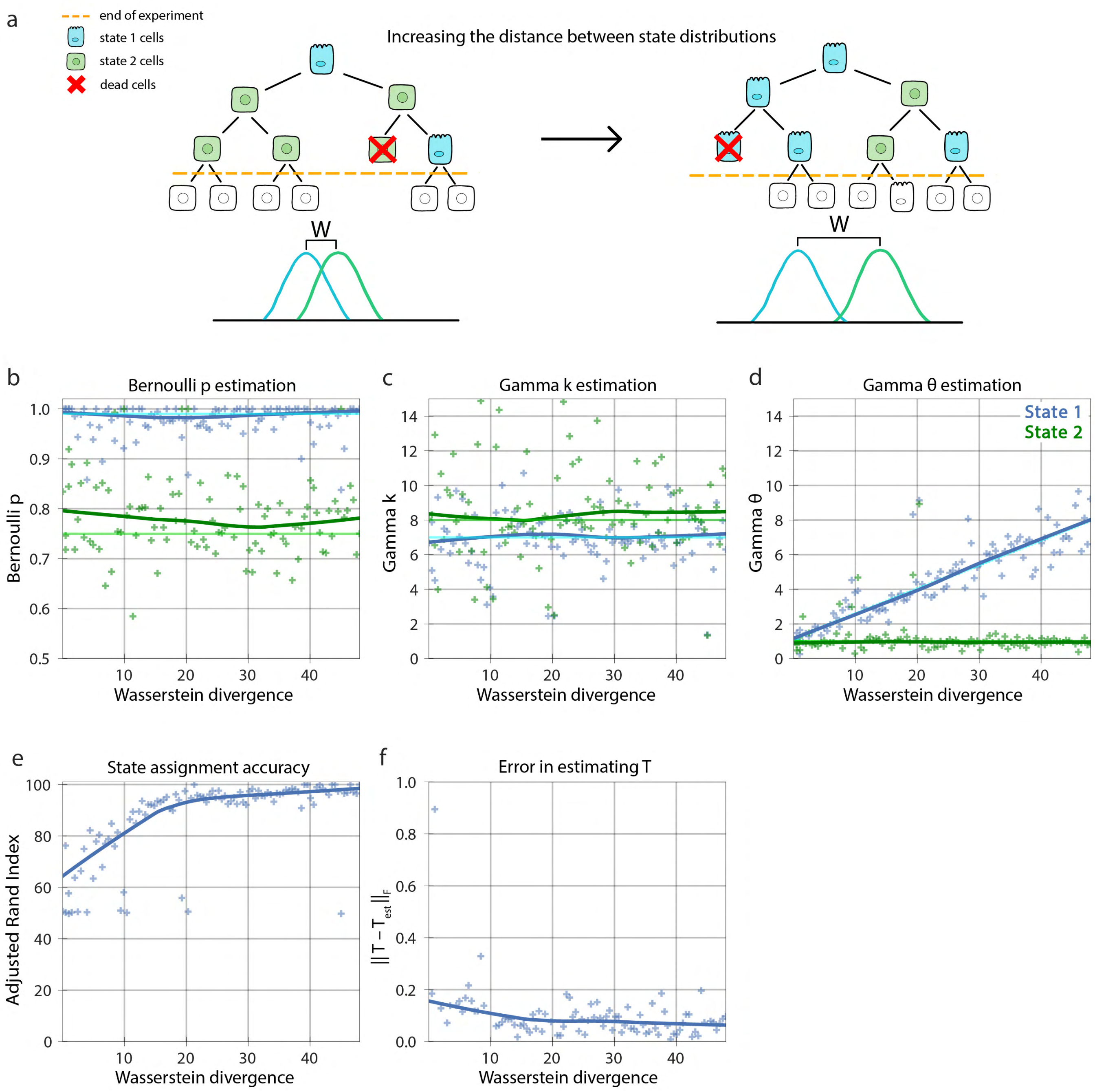


**Supplementary Figure 7: Change in model performance when varying state distribution similarity for a censored population of lineages in a synthetic two-state dataset. (a)** Visualization of the Wasserstein divergence increasing as the state distribution in the censored lineage varies. **(b)** The Bernoulli, **(c)** Gamma shape, and **(d)** Gamma scale parameters for states 1 and 2 as the Wasserstein divergence increases. **(e)** The state assignment accuracy as the Wasserstein divergence increases. **(f)** The error in the estimate of the transition probability matrix, T, as the Wasserstein divergence increases. In **(b-d)** the light green and blue solid lines show the true value of the parameters, and the dark green and blue solid lines show the Lowess trend of estimations.

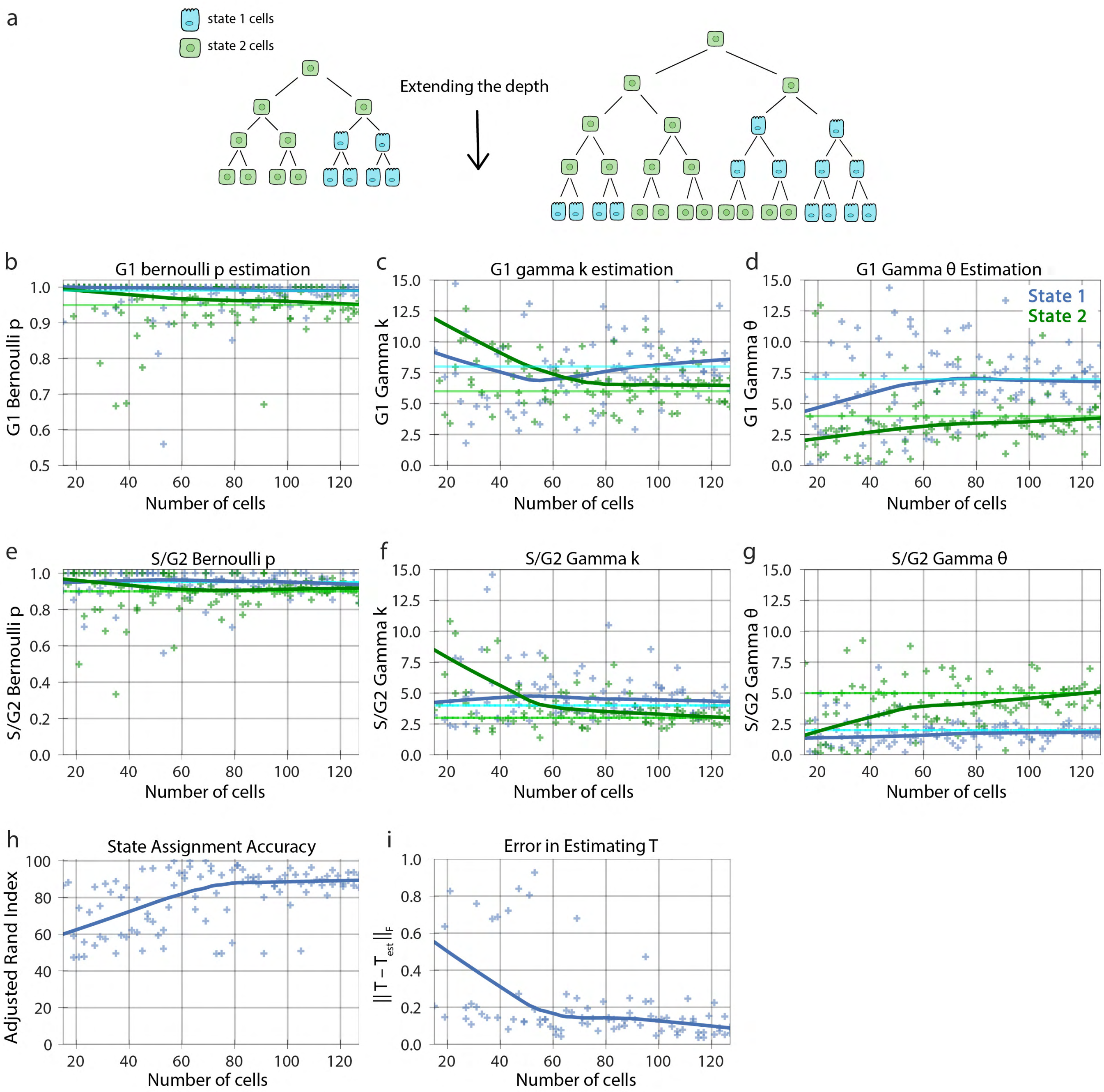


**Supplementary Figure 8: Performance of increasing cell numbers in an uncensored single lineage in a synthetic two-state dataset. (a)** Visualization of a single lineage with increasing cell number. **(b)** The G1 phase Bernoulli, **(c)** Gamma shape, and **(d)** Gamma scale parameters for states 1 and 2 as the number of cells increase. **(e)** The S/G2 phase Bernoulli, **(f)** Gamma shape, and **(g)** Gamma scale parameter for states 1 and 2 as the number of cells increase. **(h)** The state assignment accuracy as the number of cells increases. **(i)** The errors in the estimate of the transition probability matrix, T, as the number of cells increase. In **(b-g)** the light green and blue solid lines show the true value of the parameters, and the dark green and blue solid lines show the Lowess trend of estimations.

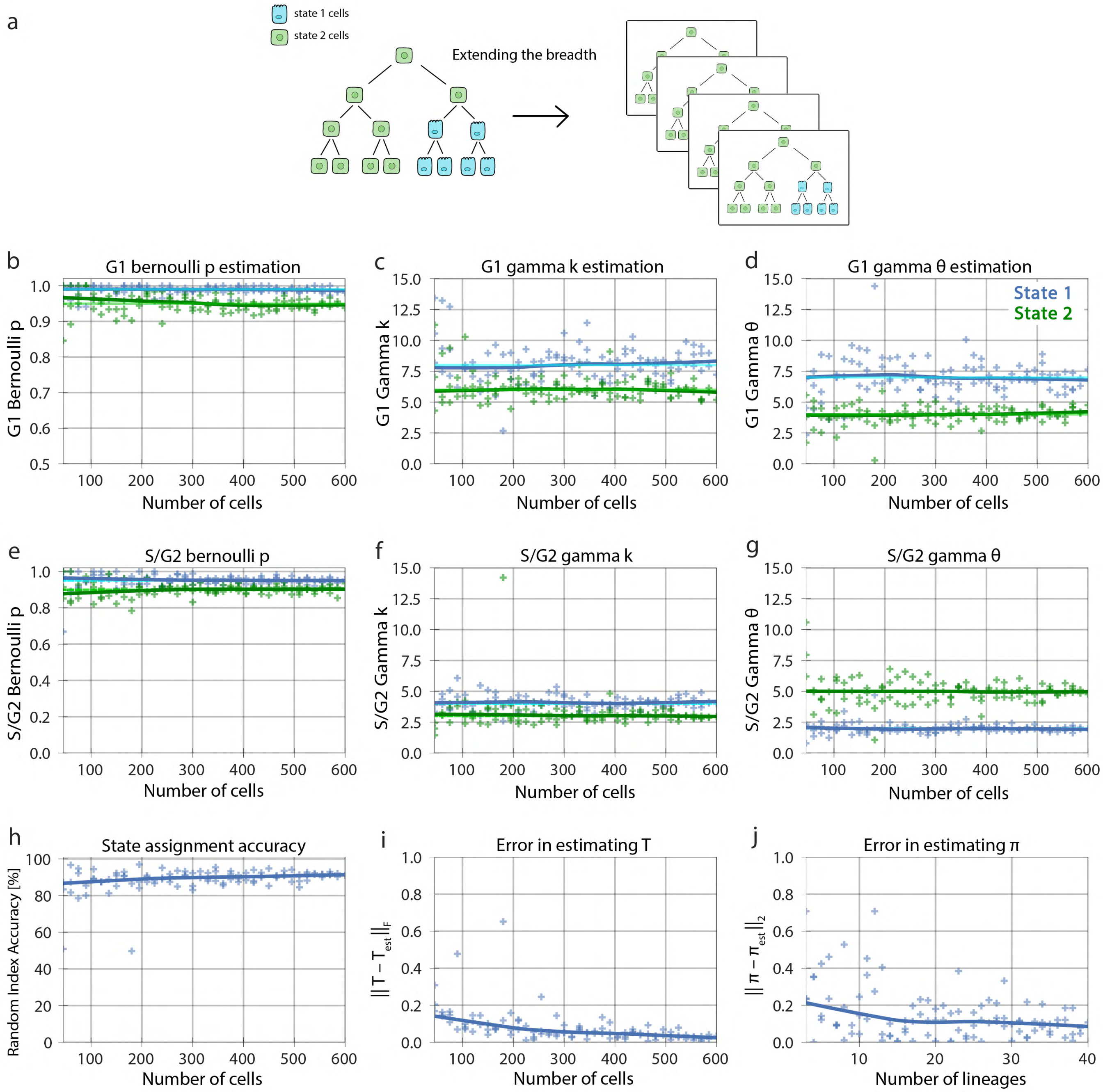


**Supplementary Figure 9: Performance of increasing lineage numbers in an uncensored population in a synthetic two-state dataset. (a)** Visualization of the number of uncensored lineages within a population increasing. **(b)** The G1 phase Bernoulli, **(c)** Gamma shape, and **(d)** Gamma scale parameters for states 1 and 2 as the number of cells increase. **(e)** The S/G2 phase Bernoulli, **(f)** Gamma shape, and **(g)** Gamma scale parameters for states 1 and 2 as the number of cells increase. **(h)** The state assignment accuracy as the number of cells increases. **(i)** The error in the estimate of the transition probability matrix, T, as the number of cells increase. **(j)** The errors in the estimate of the initial probability matrix, π, as the number of lineages increase. In **(b-g)** the light green and blue solid lines show the true value of the parameters, and the dark green and blue solid lines show the Lowess trend of estimations.

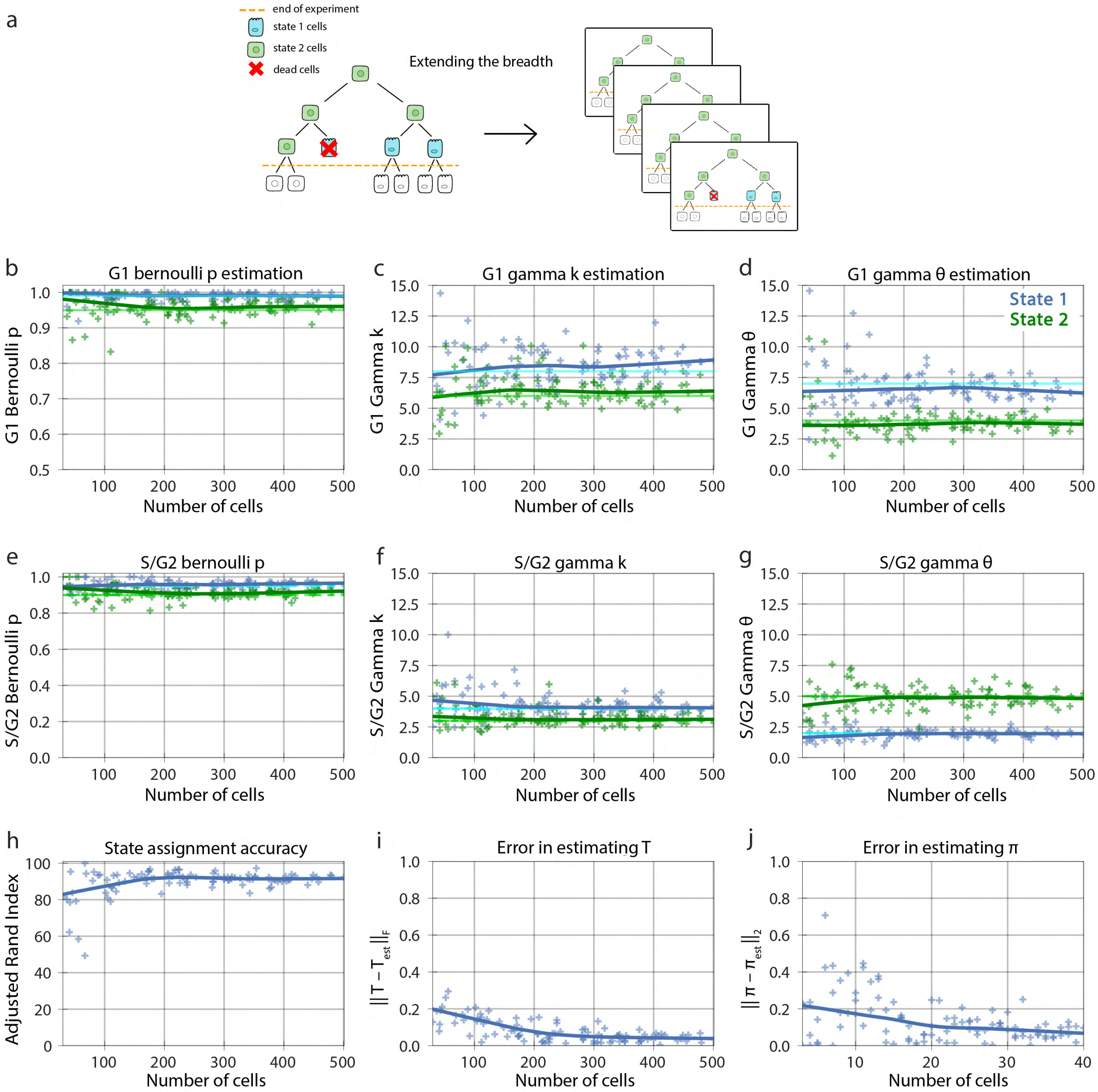


**Supplementary Figure 10: Performance of increasing lineage numbers in a censored population in a synthetic two-state dataset. (a)** Visualization of the number of censored lineages within a population increasing. **(b)** The G1 phase Bernoulli, **(c)** Gamma shape, and **(d)** Gamma scale parameters for states 1 and 2 as the number of cells increase. **(e)** The S/G2 phase Bernoulli, **(f)** Gamma shape, and **(g)** Gamma scale parameter for states 1 and 2 as the number of cells increase. **(h)** The state assignment accuracy as the number of cells increases. **(i)** The error in the estimate of the transition probability matrix, T, as the number of cells increase. **(j)** The error in the estimate of the initial probability matrix, π, as the number of lineages increase. In (b-g) the light green and blue solid lines show the true value of the parameters, and the dark green and blue solid lines show the Lowess trend of estimations.

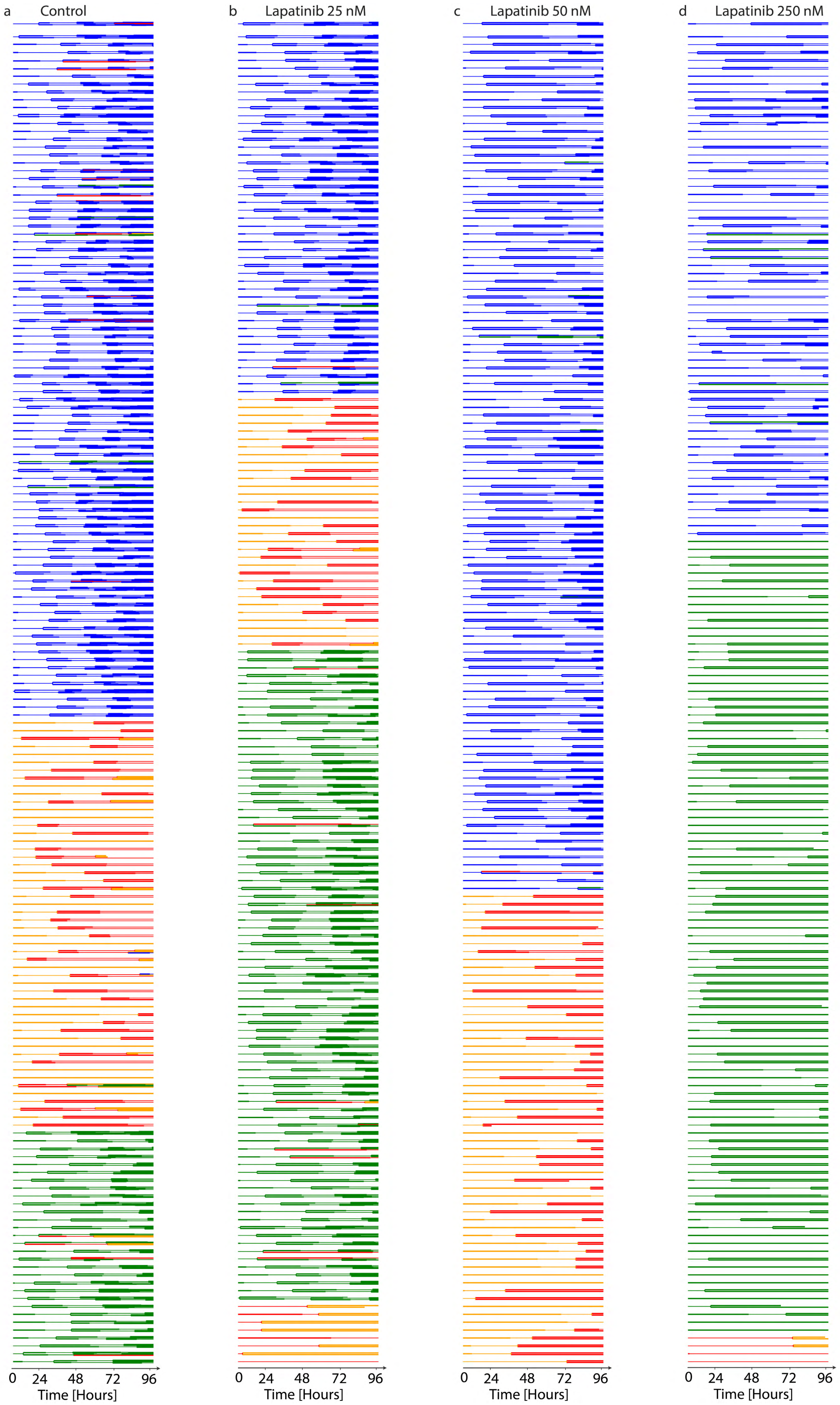
a Control
b Lapatinib 25 nM
c Lapatinib 50 nM
d Lapatinib 250 nM
0 24 48 72 96
Time [Hours]
0 24 48 72 96
Time [Hours]
0 24 48 72 96
Time [Hours]
0 24 48 72 96
Time [Hours]

**Supplementary Figure 11: The single cell data after fitting and state assignment for lapatinib-treated lineages. (a)** Control, **(b)** 25 nM, **(c)** 50 nM, and **(d)** 250 nM lapatinib treatment. Each line represents a cell, and the length of the line represents the cell's lifetime. G1 and S/G2 phase durations are depicted by thick and thin lines, respectively. Termination and branching indicate cell death and division, respectively. Different colors show the state of each cell. Blue: state 1, orange: state 2, green: state 3, red: state 4, purple: state 5, and olive: state 6.

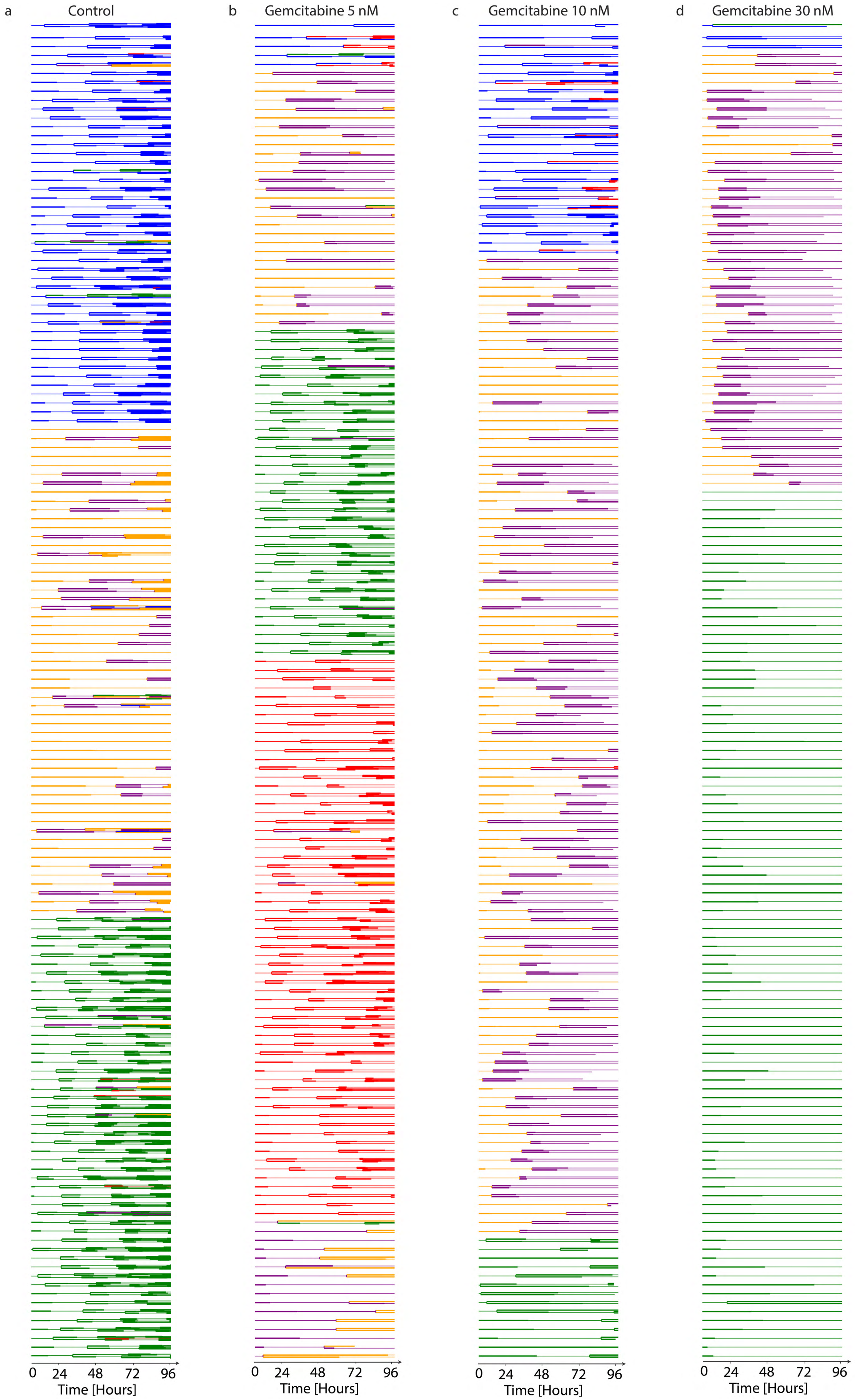
a
Control
b
Gemcitabine 5 nM
c
Gemcitabine 10 nM
d
Gemcitabine 30 nM
0 24 48 72 96
Time [Hours]

**Supplementary Figure 12: The single cell data after fitting and state assignment for gemcitabine-treated lineages. (a)** Control, **(b)** 5 nM, **(c)** 10 nM, and **(d)** 30 nM gemcitabine treatment. Each line is a cell, and the length of the line represents the cell's lifetime. G1 and S/G2 phase durations are depicted by thick and thin lines, respectively. Termination and branching indicate cell death and division, respectively. Different colors show the state of each cell. Blue: state 1, orange: state 2, green: state 3, red: state 4, and purple: state 5.

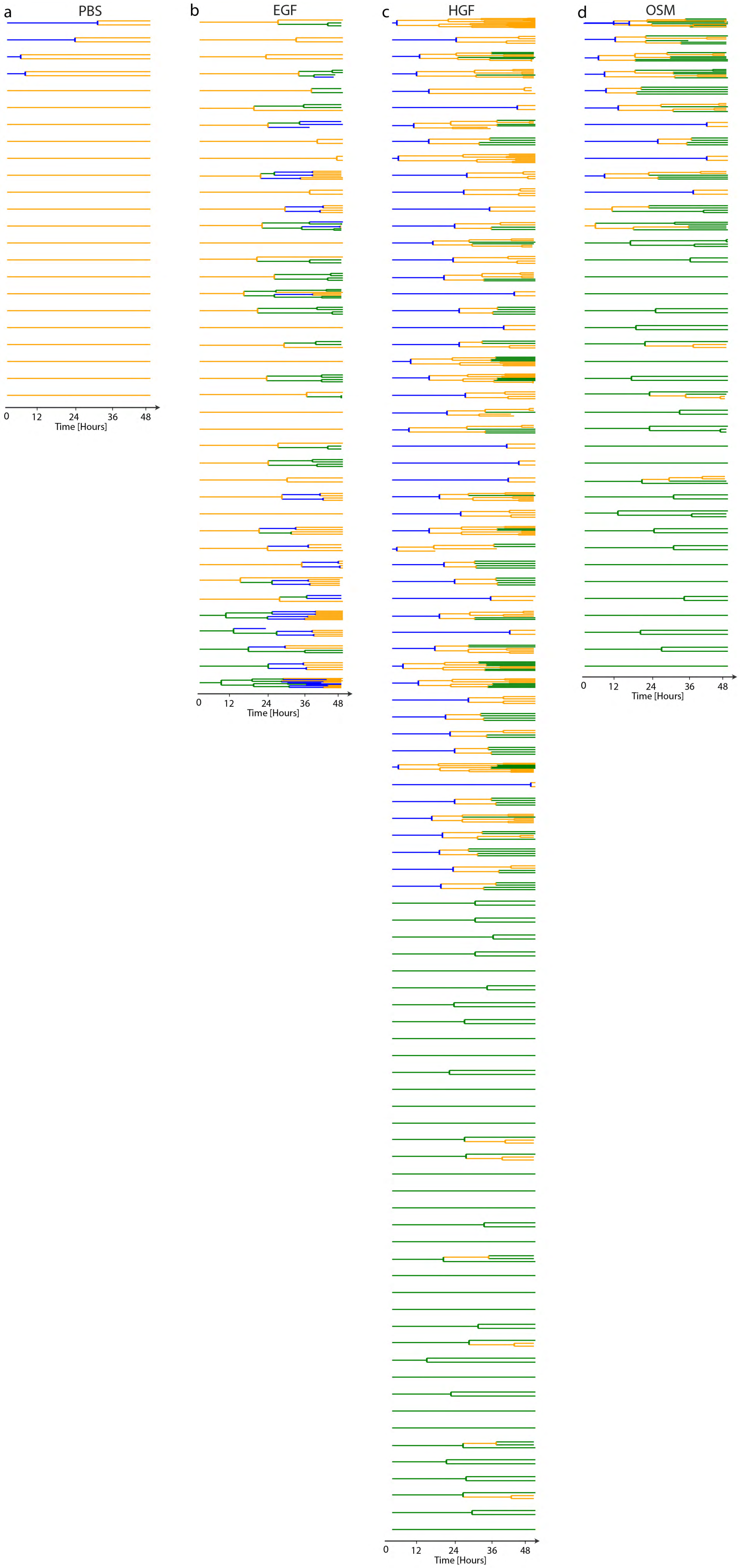
a
PBS
0
12
24
36
48
Time [Hours]
b
EGF
0
12
24
36
48
Time [Hours]
c
HGF
0
12
24
36
48
Time [Hours]
d
OSM
0
12
24
36
48
Time [Hours]

**Supplementary Figure 13: The single cell data after fitting and state assignment for growth factor treated MCF10A lineages. (a)** PBS, **(b)** EGF, **(c)** HGF, and **(d)** OSM treatments. Each line represents a cell, and the length of the line represents the cell's lifetime. Termination and branching indicate cell death and division, respectively. Different colors show the state of each cell. Blue: state 1, orange: state 2, and green: state 3.

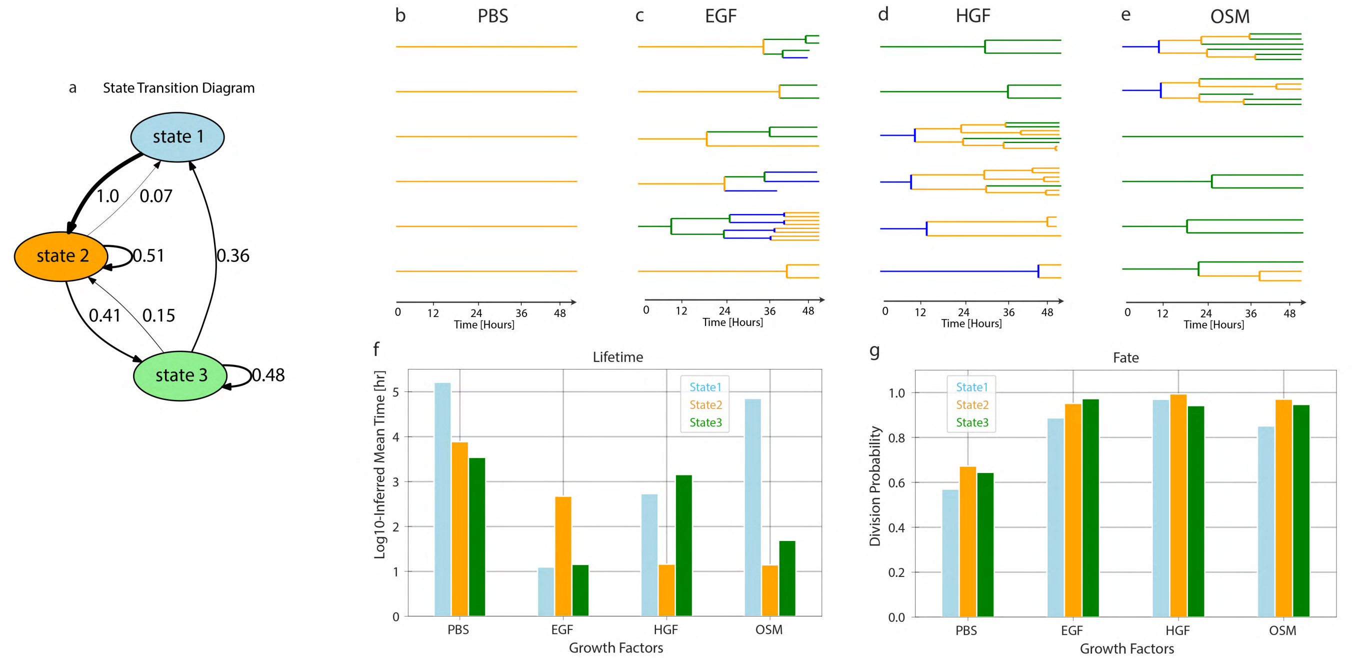


**Supplementary Figure 14: State-specific emissions of the growth factor treated MCF10A population. (a)** State transition graph showing the probability of state transitions among the states. Transitions with less than 0.03 probability have been removed. **(b-e)** A sample of lineage trees after fitting the model and state assignment (PBS, EGF, HGF, and OSM). **(f-g)** The inferred log10 mean time of cell cycle durations for different conditions at different states. **(h-i)** The Bernoulli parameter, which is the probability of dividing versus dying for different conditions and states.

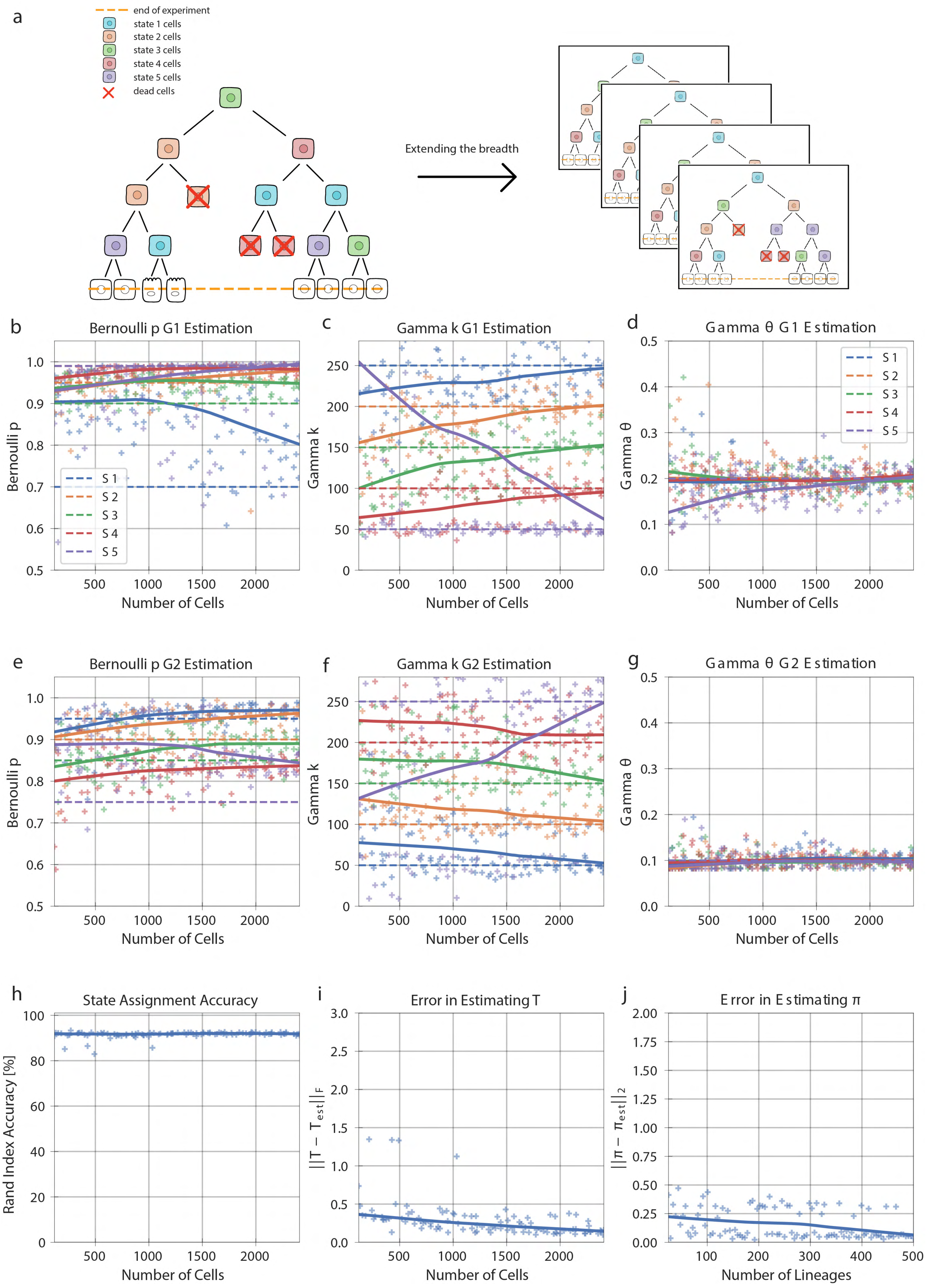
a
end of experiment
state 1 cells
state 2 cells
state 3 cells
state 4 cells
state 5 cells
dead cells
Extending the breadth
b
Bernoulli p G1 Estimation
Bernoulli p
Number of Cells
c
Gamma k G1 Estimation
Gamma k
Number of Cells
d
Gamma θ G1 Estimation
Gamma θ
Number of Cells
S 1
S 2
S 3
S 4
S 5
e
Bernoulli p G2 Estimation
Bernoulli p
Number of Cells
f
Gamma k G2 Estimation
Gamma k
Number of Cells
g
Gamma θ G2 Estimation
Gamma θ
Number of Cells
h
State Assignment Accuracy
Rand Index Accuracy [%]
Number of Cells
i
Error in Estimating T
||T − T_est||_F
Number of Cells
j
Error in Estimating π
||π − π_est||_2
Number of Lineages

**Supplementary Figure S15: Performance of increasing lineage numbers in a censored population in a synthetic five-state dataset.** **(a)** Visualization of the number of censored lineages within a population increasing. **(b)** The G1 phase Bernoulli, **(c)** Gamma shape, and **(d)** Gamma scale parameters for states 1, 2, …, 5 as the number of cells increase. **(e)** The S/G2 phase Bernoulli, **(f)** Gamma shape, and **(g)** Gamma scale parameter for states 1, 2, …, 5 as the number of cells increase. **(h)** The state assignment accuracy as the number of cells increases. **(i)** The error in the estimate of the transition probability matrix, T, as the number of cells increase. **(j)** The error in the estimate of the initial probability matrix, $\pi$, as the number of lineages increase. In **(b-g)** the light green and blue solid lines show the true value of the parameters, and the dark green and blue solid lines show the Lowess trend of estimations.

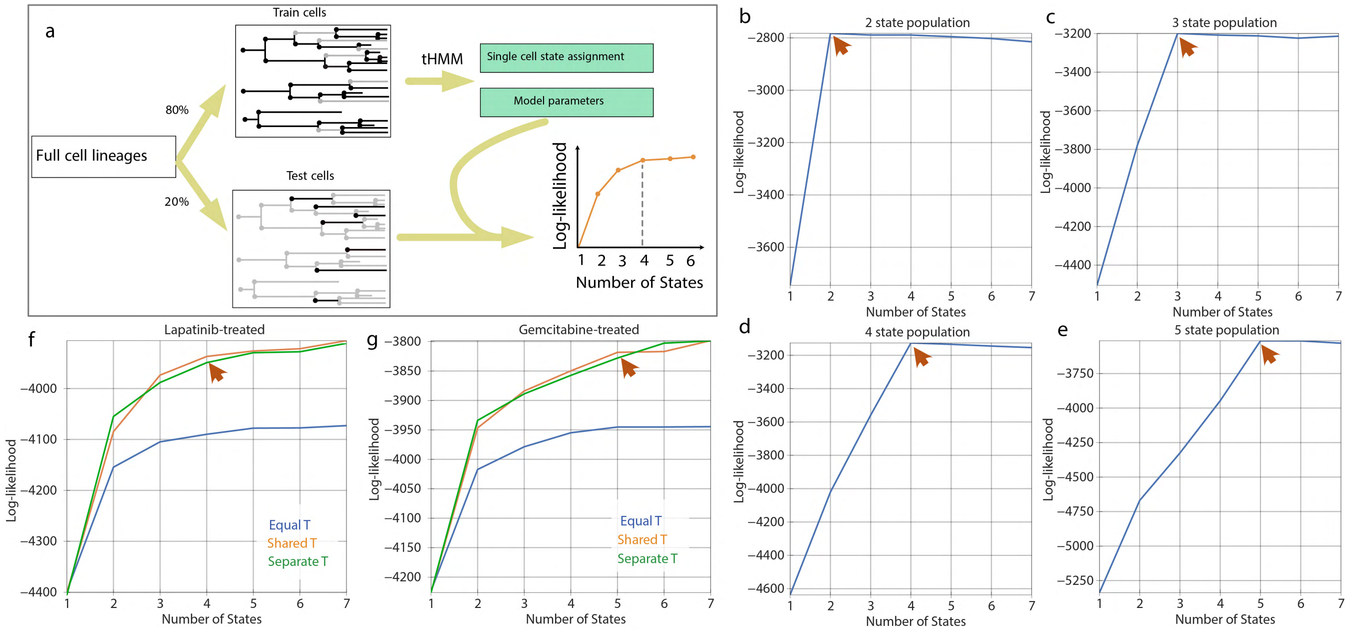


**Supplementary Figure S16: Performance of increasing lineage numbers in a censored population in a synthetic five-state dataset. (a)** Visualization of the cross-validation process. 80% of cells are randomly selected as the training set, and the remaining cells serve as the test set. The log likelihood of the observations of the test given the trained model will determine the optimum number of states. **(b–d)** The log likelihood plot for a 2, 3, 4, and 5 state synthetic model using the cross-validation scheme. **(f–g)** The log likelihood from the cross-validation approach for lapatinib and gemcitabine-treated AU565 cells. “Equal T” refers to the scenario where all transitions to and from each state are equal, simulating the absence of inheritance, “Shared T” refers to the scenario where we estimate a transition matrix that is shared between all concentrations, and “Separate T” simulates the scenario where each concentration has a separate transition matrix.
